# Sub-threshold resonance organizes activity and optimizes learning in neural networks

**DOI:** 10.1101/195040

**Authors:** James P. Roach, Aleksandra Pidde, Eitan Katz, Jiaxing Wu, Nicolette Ognjanovski, Sara J. Aton, Michal R. Zochowski

## Abstract

Network oscillations across and within brain areas are critical for learning and performance in memory tasks. While a large amount of work has focused on the generation of neural oscillations, their effects on neuronal populations’ spiking activity and information encoding is less known. Here, we use computational modeling and *in vivo* recording to demonstrate that a shift in sub-threshold resonance can interact with oscillating input to ensure that networks of neurons properly encode new information represented in external inputs to the weights of recurrent synaptic connections. Using a neuronal network model, we find that due to an input-current dependent shift in their resonance response, individual neurons in a network will arrange their phases of firing to represent varying strengths of their respective inputs. As networks encode information, neurons fire more synchronously, and this effect limits the extent to which further “learning” (in the form of changes in synaptic strength) can occur. We also demonstrate that sequential patterns of neuronal firing can be accurately stored in the network; these sequences are later reproduced without external input (in the context of sub-threshold oscillations) in both the forward and reverse directions (as has been observed following learning *in vivo*). To test whether a similar mechanism could act *in vivo*, we show that periodic stimulation of hippocampal neurons coordinates network activity and functional connectivity in a frequency-dependent manner. We conclude that sub-threshold resonance provides a plausible network-level mechanism to accurately encode and retrieve information without over-strengthening connections between neurons.

## I. Introduction

Oscillations in local field potential (LFP) largely reflect coherent post-synaptic potentials among neurons [1]. These oscillations, or rhythms, are behaviorally relevant, and their features are highly predictive of cognitive processes in underlying neural networks [2, 1, 3, 4]. However, it is unknown whether rhythms are byproducts or critical components of neural computation.

Neurons display complex behavior in response to oscillatory input. Many neuronal subtypes display sub-threshold membrane resonance - which manifests as enhanced membrane voltage responses to periodic inputs in narrow frequency bands [5, 6, 7]. Critically, the preferred frequencies of neurons have been observed shift in response to depolarizing and hyperpolarizing inputs [8, 9, 10]. This suggests that in addition to simply integrating inputs to fire a spike, neurons are biophysically suited to perform time-dependent computations and selectively filter inputs based on their periodicity.

The theta (4-10 Hz) rhythm is one of the major oscillations present in mammalian brain networks [3]. Within the hippocampus, the theta rhythm plays a central role in the function of place cells, which encode spatial and contextual information [11, 12]. Place cells show several interesting firing features with respect to the theta rhythm. First, these cells have a changing firing phase relationship relative to hippocampal theta (theta phrase precession) that changes with the location of an animal in their environment [13, 14, 15]. Second, following sequential place cell activation in the context of behavior (e.g., exploration), these same sequences of place cell firing are replayed during theta oscillations, in either the forward or reverse direction [16, 17, 18, 19]. While the idea that theta plays a role in the function of the hippocampal network is widely accepted, the underlying mechanisms for phase precession and replay are still largely unknown.

Networks of neurons that display sub-threshold resonance shifts (i.e. the firing response to oscillating input changes as a neuron is depolarized) show enhanced pattern formation and separation when oscillating inputs are present [20, 21]. Here we show that resonating networks have a firing pattern that is highly beneficial for both the encoding and retrieval of information (i.e., a pattern of external inputs). Using conductance-based model neurons, which display sub-threshold resonance, we show that networks will organize the firing of neurons around an oscillation in a manner that represents an external input. When synapses are able to evolve via a spike-timing dependent plasticity (STDP) rule, an input will be reliably encoded within the synaptic weights of a network. This leads to the subsequent reproduction of the input-induced firing pattern in the absence of the external pattern for both static and temporally dynamic inputs. We also show that sub-threshold resonance provides a network-level mechanism both for theta phase precession and for forward and reverse replay. Finally, we find that sub-threshold periodic input induces stable, highly organized functional connectivity over the theta band, in both simulated and *in vivo* networks. This work demonstrates that sub-threshold resonance organizes neuronal firing phase with respect to network rhythms, and thereby facilitates the encoding and retrieval of information.

## II. Methods

All data and code generated for this manuscript will be accessible through XXXX data archival/ public database service.

### Neuronal network model

We use a network model that is composed of *N* = 300 (or *N* = 1000 for the data in figure 5) excitatory neurons. Neuronal dynamics were of Hodgkin Huxley type and governed by the current balance equation:

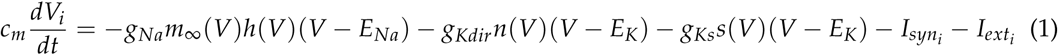

The gating variables *h*, *n*, and *s* were of the form *dx*/*dt* = (*x*_∞_(*V*) − *x*)/*τ*_*x*_(*V*). The slow potassium conductance, whose maximum value is *g*_*Ks*_, is largely responsible for the sub-threshold resonance displayed by this neuron model and its value was set to 1.5 *mS*/*cm*^2^. Additional details of the neuronal dynamics can be found in [22].

Synaptic input was modeled as a double exponential conductance pulse with the dynamics:

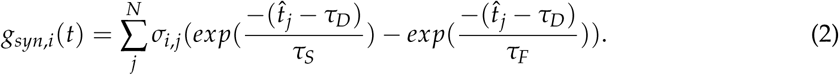

The decay constants, *τ*_*S*_ and *τ*_*F*_, were set to 250.0 and 0.3 ms respectively. The synaptic delay constant, *τ*_*D*_, was set to 0.08 ms and 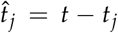 where *t*_*j*_ is the time of the last spike of the presynaptic neuron *j*. Total synaptic current to a neuron was defined as *I*_*syn*,*i*_ = *g*_*syn*,*i*_(*V*_*i*_–*E*_*syn*_) where *E*_*syn*_ is 0 mV. Networks had a connectivity rate of 6%. The connectivity scheme was small world and achieved through the Watts-Strogatz method with a rewiring probability of 0.2 [23].

Network activity was driven in two distinct ways by applying a pattern to *I*_*ext*,*i*_. For the data in figure 5 a 4 Hz oscillating current with an amplitude of 334 *nA*/*cm*^2^ was applied on top of a slowly varying activation current defined by the modified gaussian function:

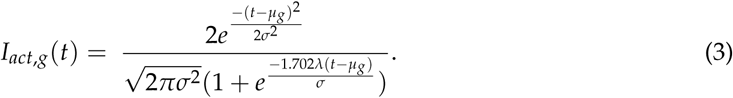

where *g* is the group to which a neuron is assigned (one of five groups), *μ*_*g*_ is the time of maximum activation of that group, *σ* = 4000 ms the width of the activation function, and *λ* = 8.0 is the skewness parameter. This leads to an activation time course that slowly grows to 227 *nA*/*cm*^2^then rapidly decays to zero (5A). In all other cases networks *I*_*ext*,*i*_ takes to form of an oscillating current on top of a constant DC current.

### Stimulation and recording of hippocampal networks

All procedures were approved by the University of Michigan Institutional Animal Care and Use Committee. Pvalb-IRES-CRE mice ((B6;129P2-Pvalbtm1(cre)Arbr/J; Jackson) were crossed to B6;129S-Gt(ROSA)26Sortm32(CAG-OP4*H134R/EYFP)Hze/J mice (Jackson) to generate PV::ChR2 mice, which expressed channelrhodopsin (ChR2) in PV-expressing (PV+) interneurons. By rhythmically activating these neurons in the hippocampus with 473 nm light, principle cells within the network were received sub-threshold periodic inhibitory stimulation. For all recordings, PV::ChR2 mice ages 2-5 months (n = 4)were anesthetized with isoflurane and chlorprothixene (1 mg/kg IP). Mice were head-fixed and a 1 mm x 1 mm matrix multielectrode (250 *μm* electrode spacing; Frederick Haer Co. (FHC), Bowdoin, ME) was slowly advanced into CA1 until stable recordings (with consistent spike waveforms continuously present for at least 30 - min before baseline recording) were obtained. An optical fiber was placed adjacent to the recording array for delivery of 473 nm laser light (CrystaLaser). Power output at the fiber tip was estimated at 3-10 mW for all experiments. CA1 neurons were recorded over a 15 - min baseline period, after which PV+ interneurons were stimulated over multiple successive 15 - min periods with a range of frequencies (2 - 18 Hz, 40 ms pulses). The various stimulation frequencies were presented in a random interleaved manner, during which neuronal activity continued to be recorded. Only those neurons recorded throughout the entire experiment were included in analyses of optogenetically induced spike-field coherence and network stability changes. For in vivo data, 80 and 68 neurons, respectively, met inclusion criteria for coherence and stability analysis. This data set also appeared in [24].

### Functional network structure

Functional network structure was calculated for both simulated and recorded networks in a similar manner. The first measure was spike wave coherence which was calculated as the range of the spike-triggered average of the LFP over a window of ± 50 ms normalized by the peak amplitude of the LFP. In simulated networks the LFP was the sum of all synaptic currents. This value ranges between 0, when spikes occur randomly in the LFP oscillation, and 1, when spikes always occur at the same time. In simulated networks the LFP was the sum of synaptic currents.

The second measure of functional network structure was the stability of functional connections though time [25, 24]. The basis of functional connectivity was the average temporal proximity of spikes between neurons and given by 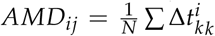 for the *i*-th to *j*-th neurons. Here 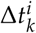 is the time difference between the *k*-th spike fired by neuron *j* and the nearest spike fired by neuron *i*. To determine whether neurons *i* and *j* are functional connected *AMD*_*ij*_ is compared to the null value given the firing rate of neuron *j* and random firing of neuron *i* by the *Z*-score 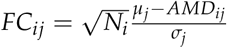. The null distribution of MD is dependent on the inter-spike intervals (ISIs)of neuron *j*. For an ISI of length *L*, the first two moments of MD are μ ^*L*^ =< *MD*^*L*^ >= *L*/4 and < (*MD*^*L*^)^2^>= *L*^2^/12. We will find an ISI of length L within a spike train of length T with a probability of *p*_*L*_ = *L*/*T*. Thus all the intervals in the spike train of neuron *j* the expected value is 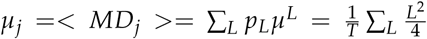. The expected standard deviation is 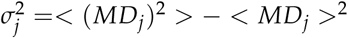 where 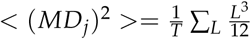. To measure the stability of inferred functional connections spiking data were separated by into non-overlapping time windows for which *FC*_*ij*_ values were aggregated into matrices *FC*_*t*_. Between adjacent time windows cosine similarity, defined by 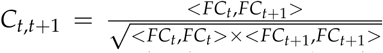, was used to quantify the change in functional network structure as a value between 0 (randomized) and 1 (no change). The stability of the functional network was quantified as the average similarity between adjacent time windows. Time windows were 2s for simulated data and 1 minute for recorded data.

## III. Results

We investigated the role that sub-threshold resonance plays in pattern and sequence learning using networks of model neurons which receive three types of input (Figure 1A). First, each cell in the network receives a unique level of external, direct current (DC) indicated by the color map. Second, the entire network receives the same oscillating input of varying frequency and magnitude. Third, individual neurons receive the summed synaptic activity defined by their presynaptic cells in the network. The weights of synapses evolve via STDP during learning phases of simulations.

**Figure 1:**
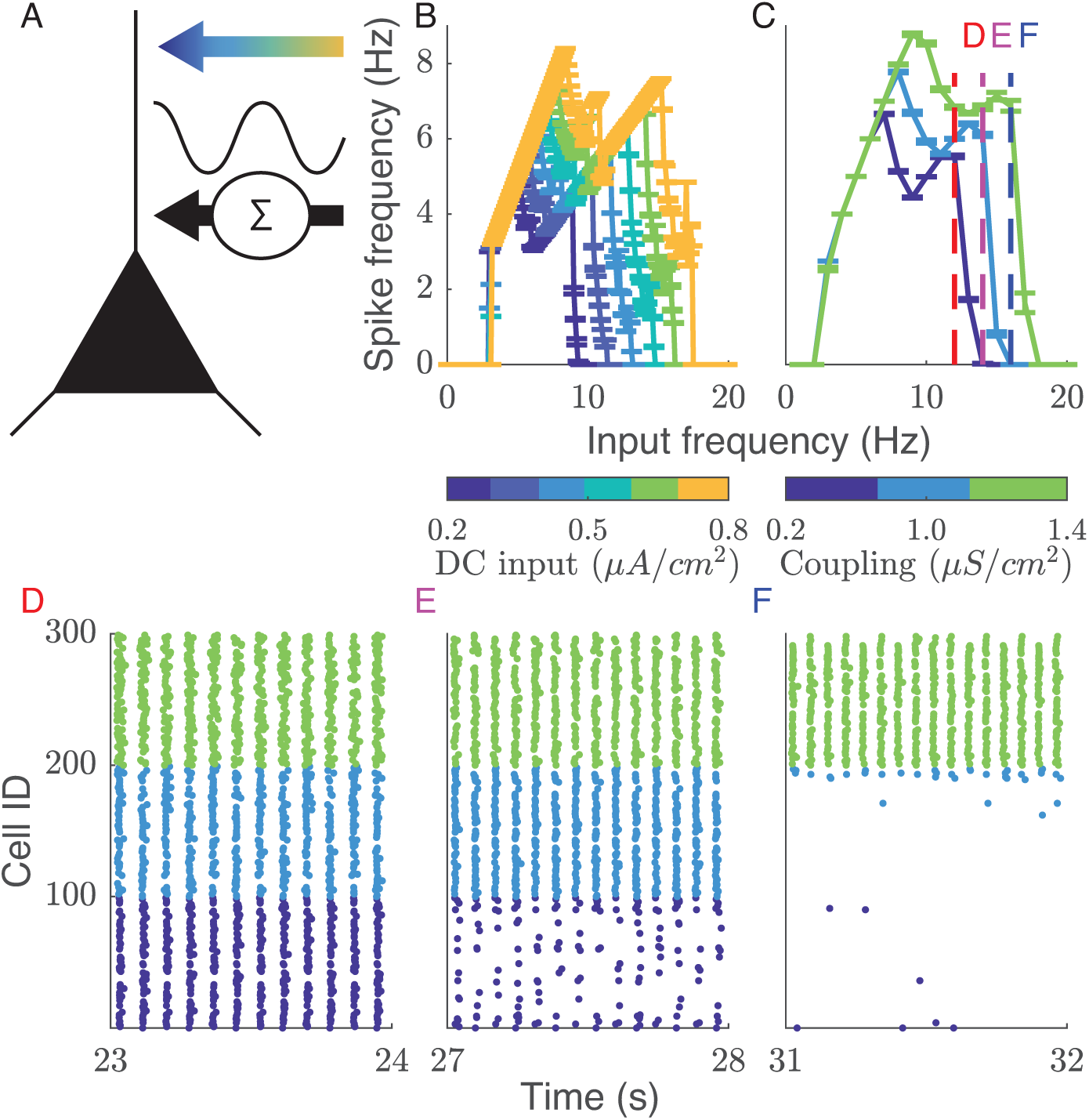
Input-dependent resonance shift allows for selectively activating subsets of neurons. (A) Model cells receive 3 types of input. External input is direct current (DC) which varies in magnitude with cell identity, represented by the color mapped arrow. All cells receive an identical oscillating input, represented by the sine wave. Additionally cells receive the synaptic inputs from neighboring cells according to network connectivity and synaptic weights. (B) The input-dependent resonance shift manifests as a broadening of the resonance curve with increasing excitation of the cells. (C) Broadening of the resonance curve also occurs for changes in synaptic weights which provides for selective activation of sunsets of cells based on synaptic coupling. Dashed lines show the frequencies corresponding to the raster plots in panels D,E,F, which show the divergent activation for frequencies between 12 and 16 Hz. Error bars = ± s.e.m.

Input dependent resonance shift allows for selective activation of subsets of neurons

The neuronal model displays an input-dependent resonance shift (Figure 1B). A neuron will respond to a wider range of oscillation frequencies if it receives a larger DC input. There are two main regimes apparent in the resonance profile, a 1:1 regime where the neuron fires one spike per cycle at low frequencies and a 1:2 regime where the neuron fires every other cycle at high input frequencies. For an oscillation of 0 Hz (i.e. in the absence of any oscillation), an additional DC current is added to the DC input so that neurons receive the same total input magnitude as when an oscillation is present. This case does not lead to neuronal spiking.

The broadening of the resonance response occurs within networks as well (Figure 1C). To show this we formed three clusters within a network with varying intra-cluster coupling (0.2, 1.0, and 1.4 *mS*/*cm*^2^), while keeping inter-cluster coupling constant. This leads to groups with high (green), moderate (light blue), and low (dark blue) synaptic input. The raster plots in Figure 1 D,E, and F show network activity at 12, 14, and 16 Hz and demonstrate how increasing the frequency of the oscillation provides for selective activation of clusters with stronger coupling.

### Networks learn patterns of external input and reproduce the reverse

To investigate the basis of learning through synaptic plasticity in this model, we had networks encode a pattern of external input (a pattern of DC inputs with varied magnitude across the network) to connections (Figure 2). We monitored phase at which the cells fire relative to the oscillations, as a function of their input magnitude. The simulations were split into five phases: prior to input pattern (red in Figure 2B), input pattern (yellow), after learning has saturated (green), and two replay periods (with and without prior patterned DC input). During the period prior to the input pattern and the replay periods all neurons received the same moderate DC input and STDP was disabled.

**Figure 2:**
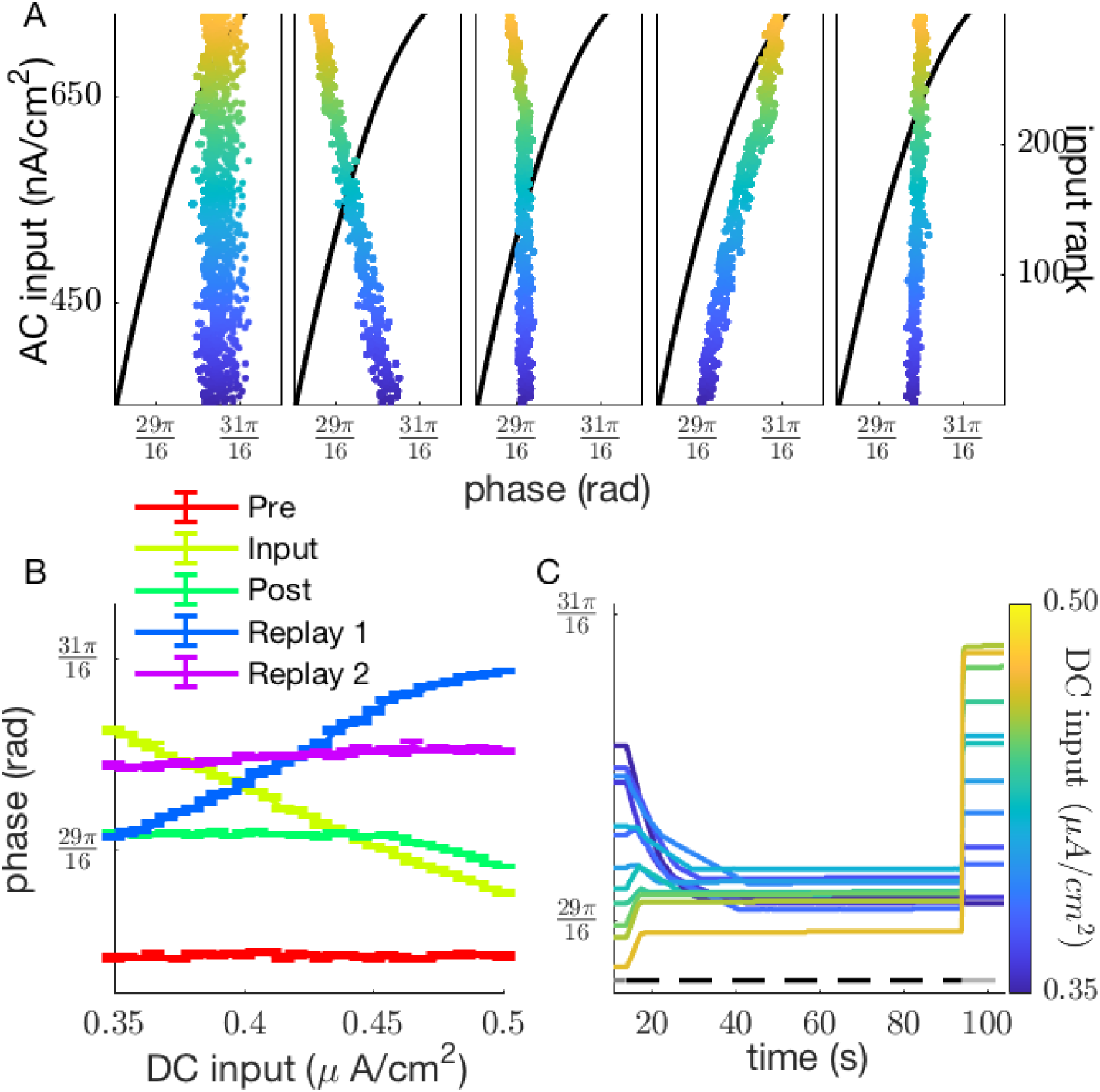
Resonating networks learn by mapping input patterns to synaptic weights. (A) Raster plots show the relationship between the phase of firing and the external input to the neuron. Black lines show the trace of the oscillating input and the color of the rasters shows the DC input to the given cell. Cells are sorted by their input rank. Sub-panels in A correspond to before DC input distribution is applied (Pre), with DC input distribution (Input), after learning has saturated (Post), after learning/ no DC distribution (Replay 1), and after a second period of learning with no DC distribution (Replay 2). (B) The relationship between firing phase and DC input varies between negatively, positively, and not correlated for different epochs of the simulation. Data are averaged over 10 cycles of the oscillation. Error bars = s.e.m. (C) Transitioning from the input-pattern depending firing phases to synchronous firing is gradual. Lines trace the firing phase of 12 neurons with varying input magnitudes across time. The horizontal bars above indicate when the external input and learning are present (dark gray -input but no learning; black - learning and input, light gray - no input and no learning).

Replay period one shows the effect of learning the input pattern and replay period two shows the effect of a second learning period where no input pattern is present.

The raster plots in Figure 2A show the evolution of firing phase across each period of the simulation. The color indicates the magnitude of input current a cell receives and cells are sorted by this value with highly activated neurons having a higher input rank. Before any input cells fire randomly over a narrow band of phases (Figure 2A far left). The input pattern leads to organized firing with highly activated cells firing at earlier phases (Figure 2A inner left). As the pattern is learned, the overall phase shifts, but cells return to firing at a uniform phase, independent of their DC input (Figure 2A center). When learning is suspended and the external input pattern is removed the network show the reverse pattern of activation (Figure 2A inner right). After a second period of learning (but with a uniform external input) the network returns to firing at a uniform phase (Figure 2A far right). The above relationships are summarized in Figure 2B as we plot relative phase of neuronal spiking as a function of their DC input magnitude. Red line (Pre) depict firing phases before DC input is activated, thus all phases remain the same. After the input is activated, but before learning, the firing cells shows strong dependence on the input magnitude with their phase decreasing with increasing magnitude of input (input; yellow line). After learning spike timing dependent plasticity is turned on the neurons again equalize their firing phases (post; green line). When the patterned input is removed (i.e. all neurons receive the same DC magnitude), the neurons activate in reverse order (blue line; replay 1). Finally, when the learning is again activated, but neurons do not received the patterned input, the phases of the neuronal firing equalize again (violet line, replay 2). Figure 2C depicts the time-course of the evolution of firing phase for 11 neurons having different DC input values. The bars above indicate timeline when input and learning are present (dark gray -input but no learning; black - learning and input, light gray - no input and no learning). The neurons initially show varied phase response to DC input, which rapidly converges during learning. This convergence is due to the universal learning rule which mimics spike timing dependent plasticity [26], namely that the connection between two neurons is strengthen if the postsynaptic cells fires after the presynaptic cell, and conversely it is weakened when the presynaptic cells fires after the postsynaptic cell. Thus, the network-wide effect of this rule is that the connections from strongly (input) driven neurons to weakly driven neurons will strengthen, these from weakly driven neurons to strongly driven neurons will weaken. After input is deactivated, after learning, the neurons activate in reverse order.

### Pattern learning saturates naturally in resonating networks

Above results indicate that the phases on neuronal activations rapidly converge during learning to minimize the phase difference between the cells. This behavior has two desired effects: 1) the synaptic changes will saturate - the synaptic efficacies will stop changing when the phases converge, and 2) the input differences between the cells are mapped onto their synaptic weights. To show these effects we presented an input (DC) pattern to network for a long time-period and tracked the time course of synaptic change. If the learning rate (the magnitude of synaptic change corresponding to Δ*t* = 0) allows, both the maximum (Figure 3A) and mean (Figure 3B) synaptic weight will saturate before the end of the simulation. Regardless of learning rate there is a large increase in synaptic change followed by a gradual decline to no change in synapse strength (Figure 3C). The time of peak synaptic change is delayed for slower learning rates. Note that the input pattern is the same for all conditions in Figure 3(A,B,C). Both the final mean synapse strength (Figure 3D black) and time it takes to saturate (Figure 3D red) depend on the range of currents in the external pattern. The time to saturation is the time it takes for the mean synaptic strength to fall within 1% of its final value.

**Figure 3:**
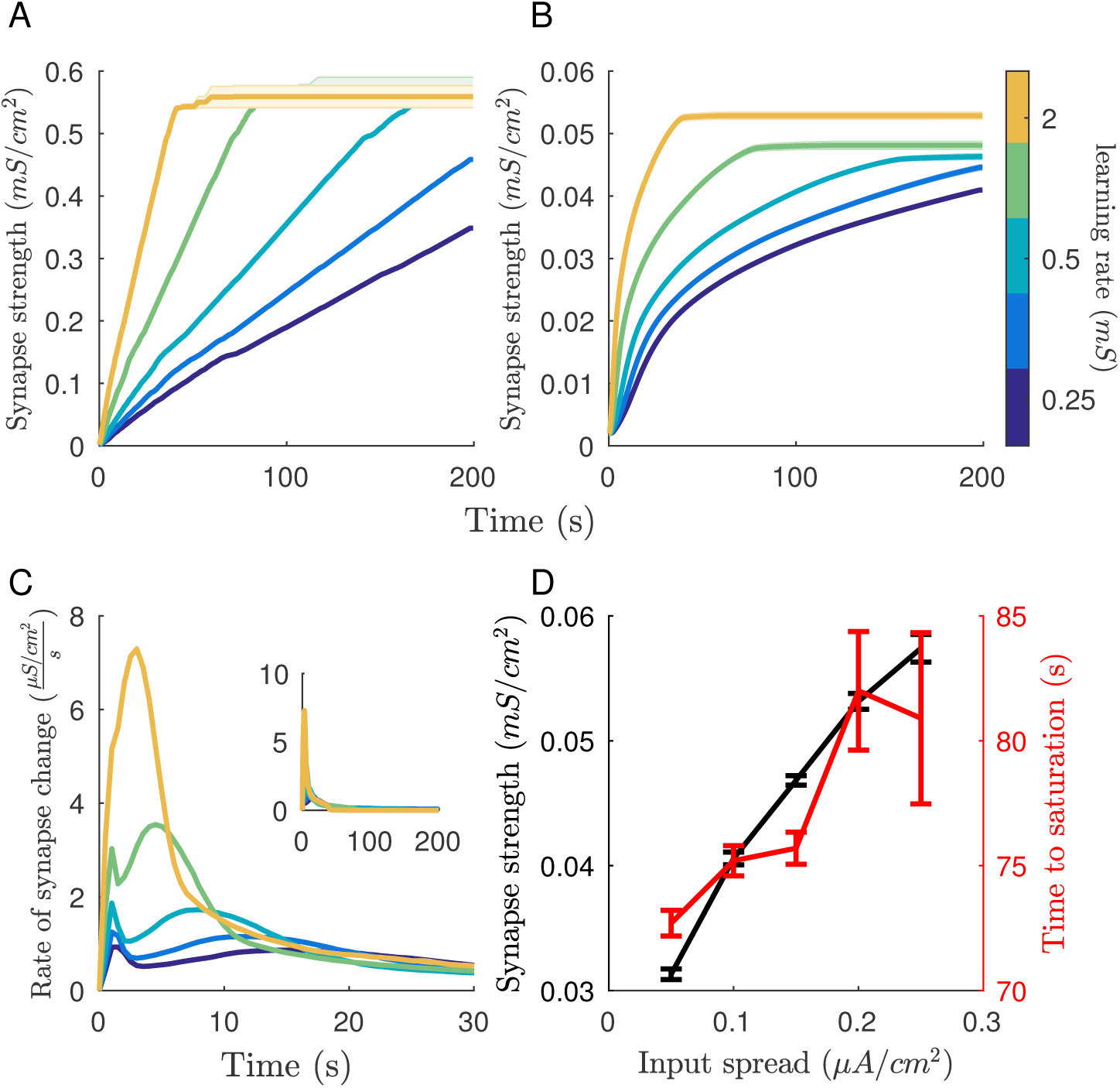
Learning saturates naturally after input pattern is completely mapped to synapses. Saturation of learning reliably occurs given that the learning rate is high enough for the given time. Both maximum (A) and mean (B) synaptic weight saturate. Line color indicates network learning rate. (C) The majority of synaptic change occurs early during the learning period then gradually decreases to zeros. Inset shows the total length of the simulation. (D) Final mean synapse strength and time until learning saturates depends on the spread of the input distribution. Error bars = ± s.e.m.

Saturation of learning occurs when the input pattern is fully mapped to the synaptic weights in the network. This is quantified in Figure 4. The mapping of the input pattern is reversed in the synaptic weights. Highly activated cells, which fire at an earlier phase, strengthen outward connections (black trace) while weakening inputs (red trace). Cells given lower external inputs do the opposite, strengthening inputs and weakening outputs. This leads to the external input pattern and the synaptic input pattern being complimentary, leading to all cell receiving the same net input.

**Figure 4:**
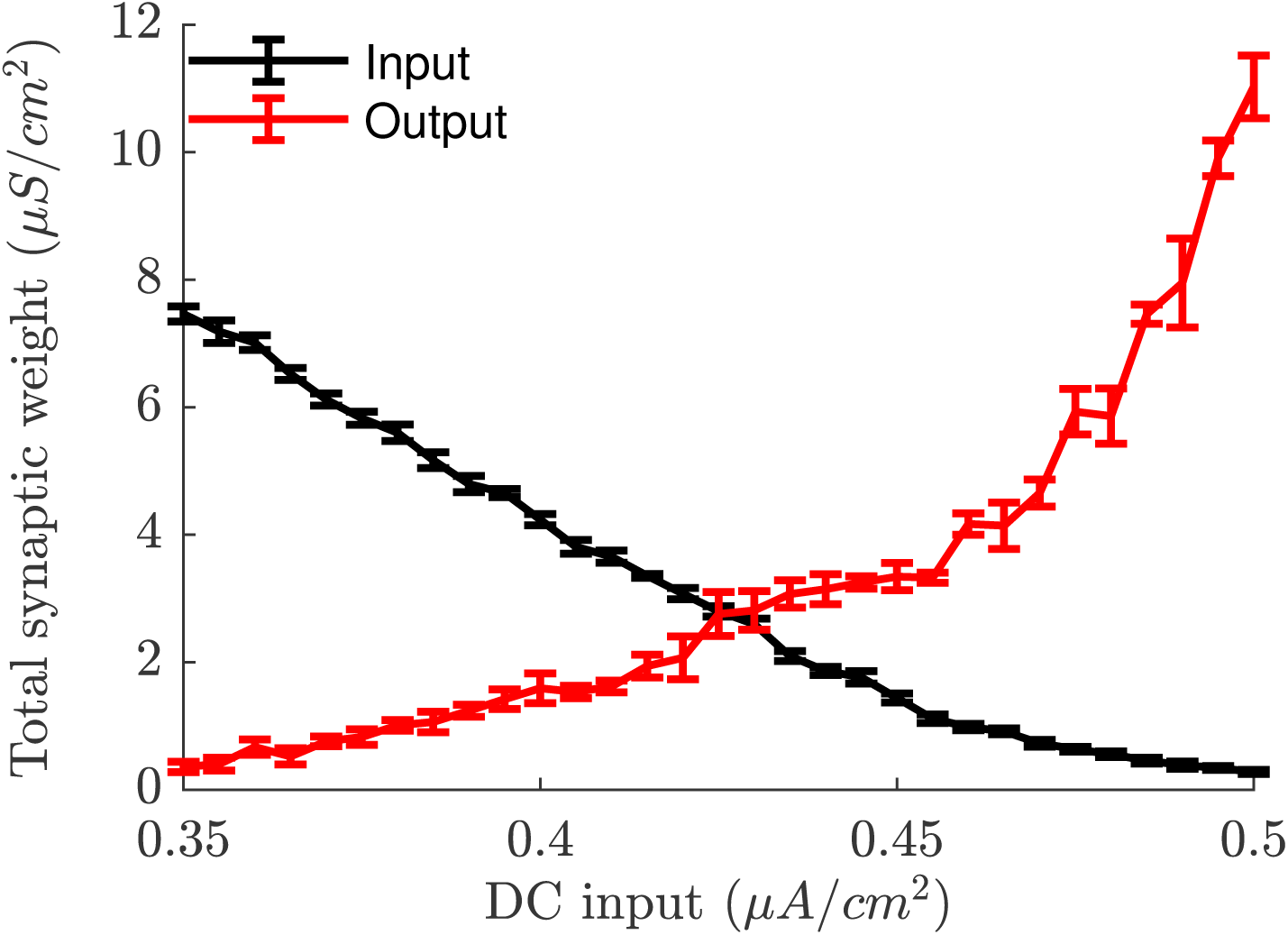
Input pattern maps to both synaptic inputs and outputs. After learning, input strength (black) is anti-correlated with input magnitude of a neuron in the pattern and output strength (red) is correlated. Error bars = ± s.e.m.

Overall, cells with the lowest DC current within the input pattern strengthen inputs more than the rest of the network while highly activated cells do the opposite. The new pattern of synaptic connectivity is complementary to the input pattern, which leads to all neurons firing at the same phase. Synchronous firing terminates learning because as spike-time differences between neurons approaches zeros there is no net synaptic change (though simplified in our model as zero synaptic change for Δ*t* < 1.5*ms*). When the external input is removed, the complementary synaptic input distribution lead to a reversal in firing order from the input pattern.

### Sub-threshold resonance facilitates sequence learning and replay

Next we investigated whether we can use sub-threshold resonance shifts to store sequential neuronal activation to model the phenomenon of sequential replay following experience [11]. Sequences were generated by delivering a slowly varying current to sequentially activate subsets of neurons (Figure 5A; solid lines), with each group resonating with the oscillating current in turn. This current is to model preferential activation of subpopulations of place cells when traverses a series of spatial locations. The asymmetry in its shape is to model the forward approach as the animal sees oncoming location. At the same time it provides input relationships between the cells to strengthen connections between the cells from prior location to the next consecutive location on the maze. The activation sequences were presented to the network 10 times during which synapses were allowed to evolve using the same learning rule as before. After this learning phase, the sequence can be reproduced in both the forward (Figure 5F) and reverse directions (Figure 5D). Both types of replay occur under different dynamical conditions. Reverse replay occurs when the whole network is depolarized to resonate with the oscillating input, but all neurons are activated to the same extent (i.e., each neuron receives the same D.C. input). This is due to the fact that more recently-activated neuronal groups receive overall larger input than groups activated previously due to asymmetry in connections. This results in earlier phase activation when the network resonates with the oscillatory current. In contrast, forward sequential replay occurs when the network is driven by external noise. The neurons which fire early in the sequence subsequently depolarize neurons at the adjacent location, making them more prone to fire. Summary data is shown in Figure 5E for reverse replay firing phase among the 5 groups. During reverse replay groups activated earlier in the sequence reliably fire at a later phase of the oscillation (Red trace). Without any learning (when STDP is off), groups generally fire at the same phase of the oscillation (black trace). During forward replay the feed-forwardness of the intergroup connections dominate. The original firing order of the groups is reproduced and early groups fire before late groups (Figure 5E; red trace).

**Figure 5:**
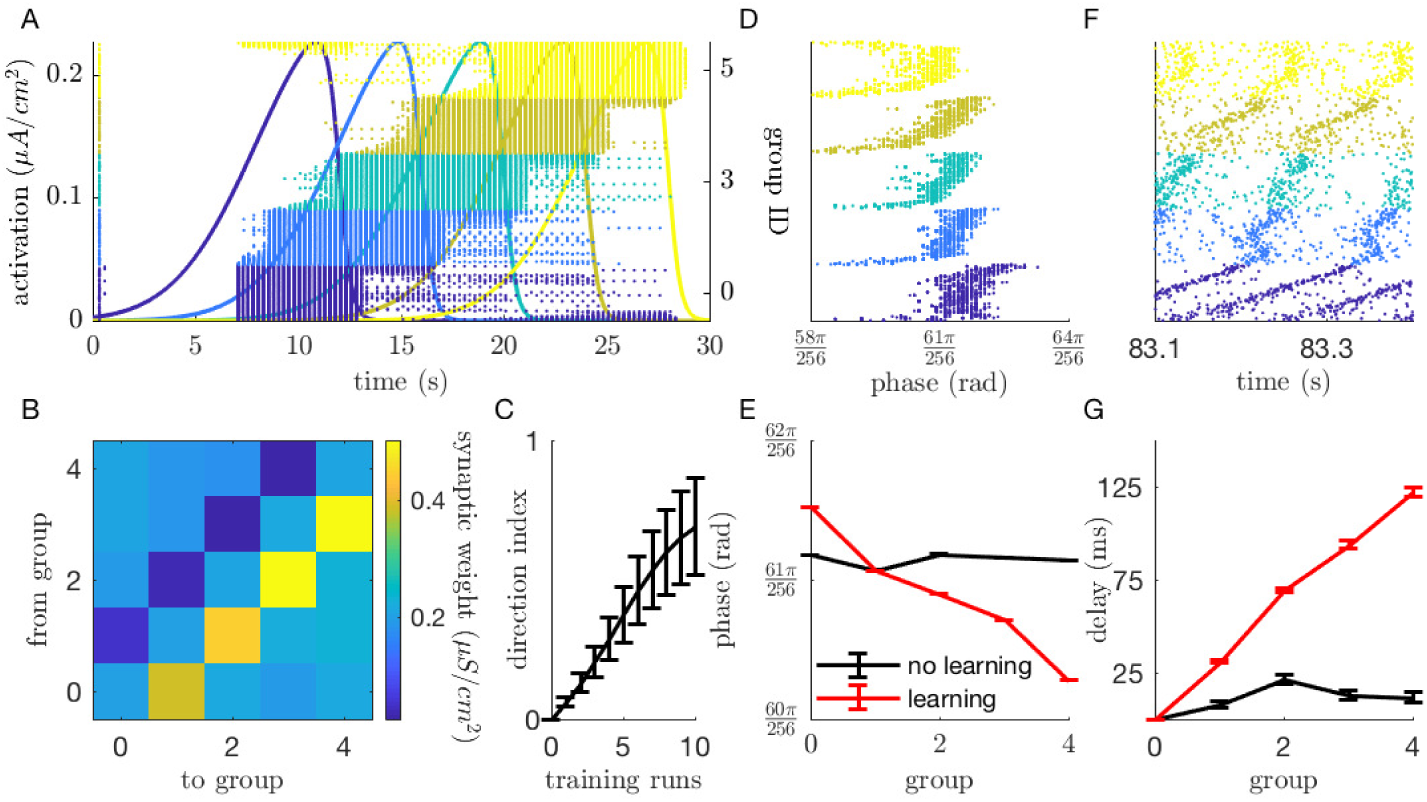
Sub-threshold resonance facilitates sequence learning. (A) Networks were trained to reproduce sequential activation of subsets of neurons. Subsets were brought into resonance with an oscillating input by adding a slowly varying current (solid lines). Spiking activity of each group is represented by the raster plots of different colors. (B) Learning induces feedforward connectivity motif within synaptic weights. Mean synaptic weights between groups show strengthened connections in the direction of the sequence and weakened connections in reverse. (C) Feedforward motif emerges steadily over training runs. (D) After learning, resonating networks display reverse reactivation of the sequence. The raster plot shows a single oscillation of network activity with time represented as phase with respect to the oscillating input. (E) Reverse reactivation is quantified for networks with (red) and without synaptic plasticity. (F) After learning non-resonating networks display reactivation of the sequence in the same order as it was presented. The raster plot shows a single oscillation of network activity with time represented as absolute simulation time. (G) The delay between group firing is quantified in with and without synaptic plasticity. Error bars = ± s.e.m.

Sequential learning leads to connections being strengthened in the same direction of the sequence (feedforward) and weakens connections in the reverse (feedback). Mean synaptic weights between groups show strengthened connections in the direction of the sequence and weakened connections in reverse (Figure 5B). This is quantified for the entire network by the direction index which is 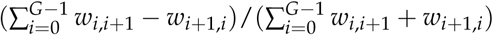, where *w*_*i*,*i*+1_ is the mean synaptic weight of connections between groups (Figure 5C).

### Functional network structure emerges in the theta band

To validate our model, we next sought to compare the behavior of simulated networks with *in vivo* networks. Information representation and subsequent encoding of information require stable spike time relationships and since sub-threshold resonance leads to stable spike-time vs phase relationships in our model we used functional connectivity as a proxy for sub-threshold resonant behavior. In networks driven by oscillatory input (a 0.3 *μA*/*cm*^2^ amplitude sine wave with a 0.3*μA*/*cm*^2^ DC offset) and background noise, oscillatory input leads to highly organized functional network structure between 4 - 10 Hz (Figure 6). We quantified functional connectivity in three ways: spike-LFP coherence, mean average minimum difference (AMD) z-score, and functional network stability. Spike-LFP coherence, which represents the reliability of the time of spikes within the LFP oscillation across the entire network, shows a noise dependent resonance effect for stimulation between 3 and 13 Hz (Figure 6A). AMD z-score and functional network stability are related measures that are based on the pairwise relationships between spike times of neurons across the network. The average significance (z-score) of AMD measures between neurons shows a narrow resonance effect between 4 and 10 Hz with a peak effect at 6 Hz which depends on the level of background noise (Figure 6B). Functional network stability, which is how similar AMD z-scores are across time and captures the stability spike-time relationships, displays a similarly narrow resonance effect between 4 - 10 Hz, but maintains a near maximal value throughout this band (Figure 6C). To validate the effects of sub-threshold stimulation on neuronal networks we optogenetically stimulated *in vivo* hippocampal networks. Rhythmic stimulation of parvalbumin-expressing (PV+) interneurons in PV::ChR2 transgenic mice was used to ensure that principle cells within the network were received sub-threshold periodic inhibitory stimulation. Rhythmic optogenetic stimulation of PV+ interneurons leads to significant increases in both spike-LFP coherence and functional network stability for frequencies between 4-10 Hz among the principle cells within the network (Figure 6D).

**Figure 6:**
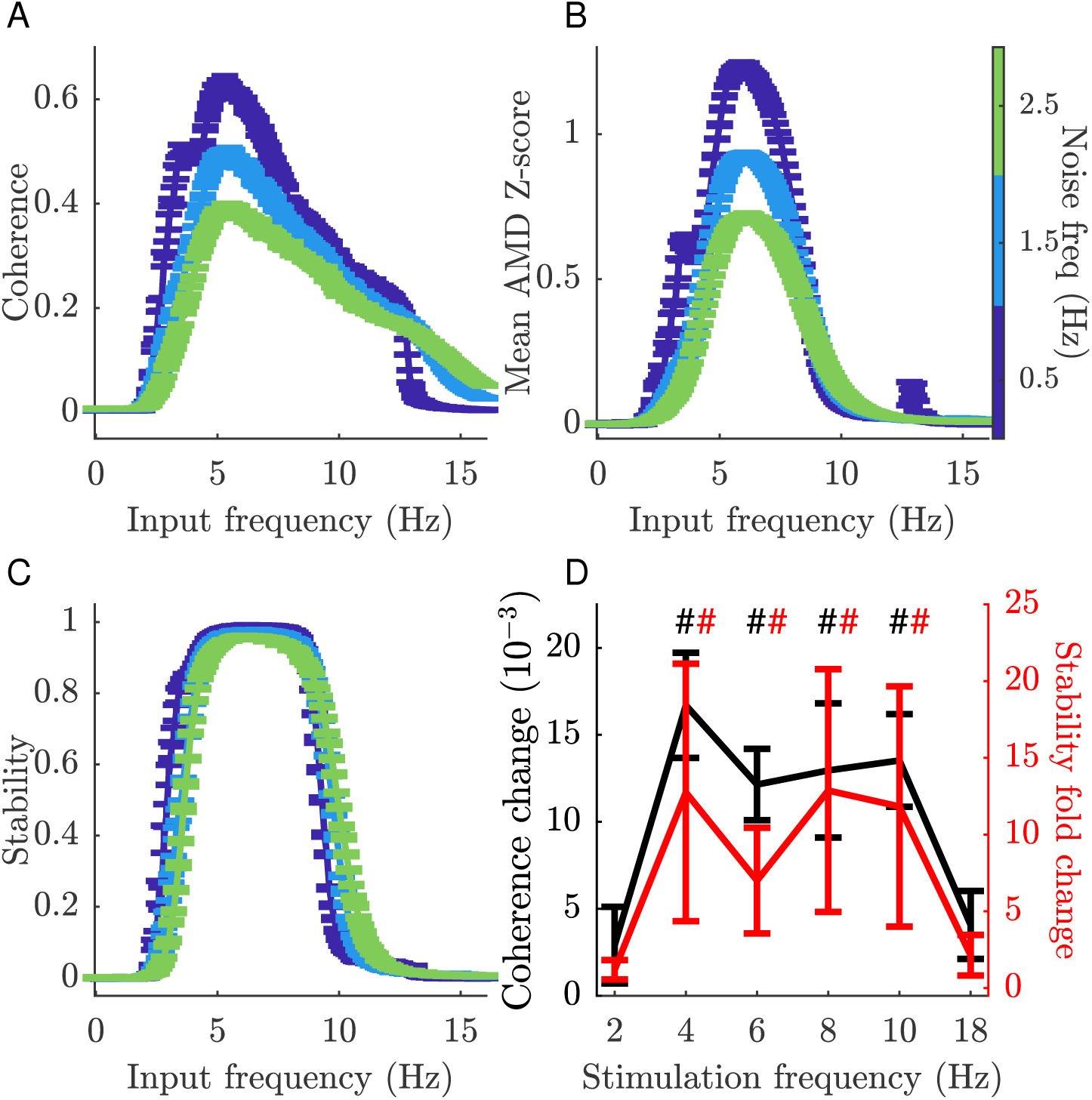
Resonating networks have organized functional structure over a narrow frequency band. Theta band resonance leads to highly organized functional network structure. In simulated networks spike-LFP coherence (A), mean AMD z-score (B), and functional network stability (C) all dramatically increase between 4-10 Hz. This effect is robust to noise, which is indicate by line color. (D) In vivo optogenetic stimulation of hippocampal PV+ neurons lead to similar increases in spike-LFP coherence and functional network stability at these frequencies. Error bars = ± s.e.m.

## IV. Discussion

We demonstrated in a biophysical model that shifting sub-threshold resonance facilitates learning of static and sequential patterns in neural networks. Our model combines sub-threshold activation of neurons by stable and oscillating cur- rents which leads to firing in a narrow frequency band. The firing rate resonance of our model neurons displays an input dependent broadening which allows for selective activation of subsets of neurons within a network. The resonance effect also leads to detailed mapping of a firing phase versus input relationship beneficial for the encoding of patterns into synaptic weights, and for the autonomous termination of learning. The resonant effect at the single neuron level leads to the emergence of highly organized spike-time relationships at the theta band which we have also shown in in vivo experiments.

The input-dependent broadening of the resonance curve in firing rate (Figure 1) allows for selective activation of subsets of neurons within a network with increasing input frequency as has been demonstrated in other computational models indicating this is a general property of neural networks with sub-threshold resonance [21]. This provides a mechanism for networks to change representations by shifting the pattern of input strengths, or alternatively, by modulation of the oscillatory input frequency. Such a mechanism would operate similarly for both externally generated (i.e. sensory input) and internal (i.e. stored representations within synapses) inputs.

The above described mechanism can simultaneously promote both forward and reverse replay of recently-learned sequences in neural networks. The reverse firing phase relationship and learning saturation seen in the external pattern simulations provide a plausible mechanism for the generation of reverse replay events in vivo (Figure 7). This mechanism relies on the fact that neurons with high input fire at early phases of oscillatory drive when in resonance. Before any synaptic change occurs the firing phase is governed by the distribution of the external inputs the cells receive. As learning progresses, neurons with the lowest external input strengthen their synaptic inputs more than the rest of the population, while highly activated neurons do the opposite. This is quantified in Figure 4. The emerging pattern of synaptic connectivity is complementary to the input pattern, which leads to all neurons firing at the same phase (i.e., in synchrony). Synchronous firing leads to no net synaptic change and thus terminates learning. As the complimentary input pattern is now represented within synaptic weights, in the absence of external input neurons fire in the reverse order.

**Figure 7:**
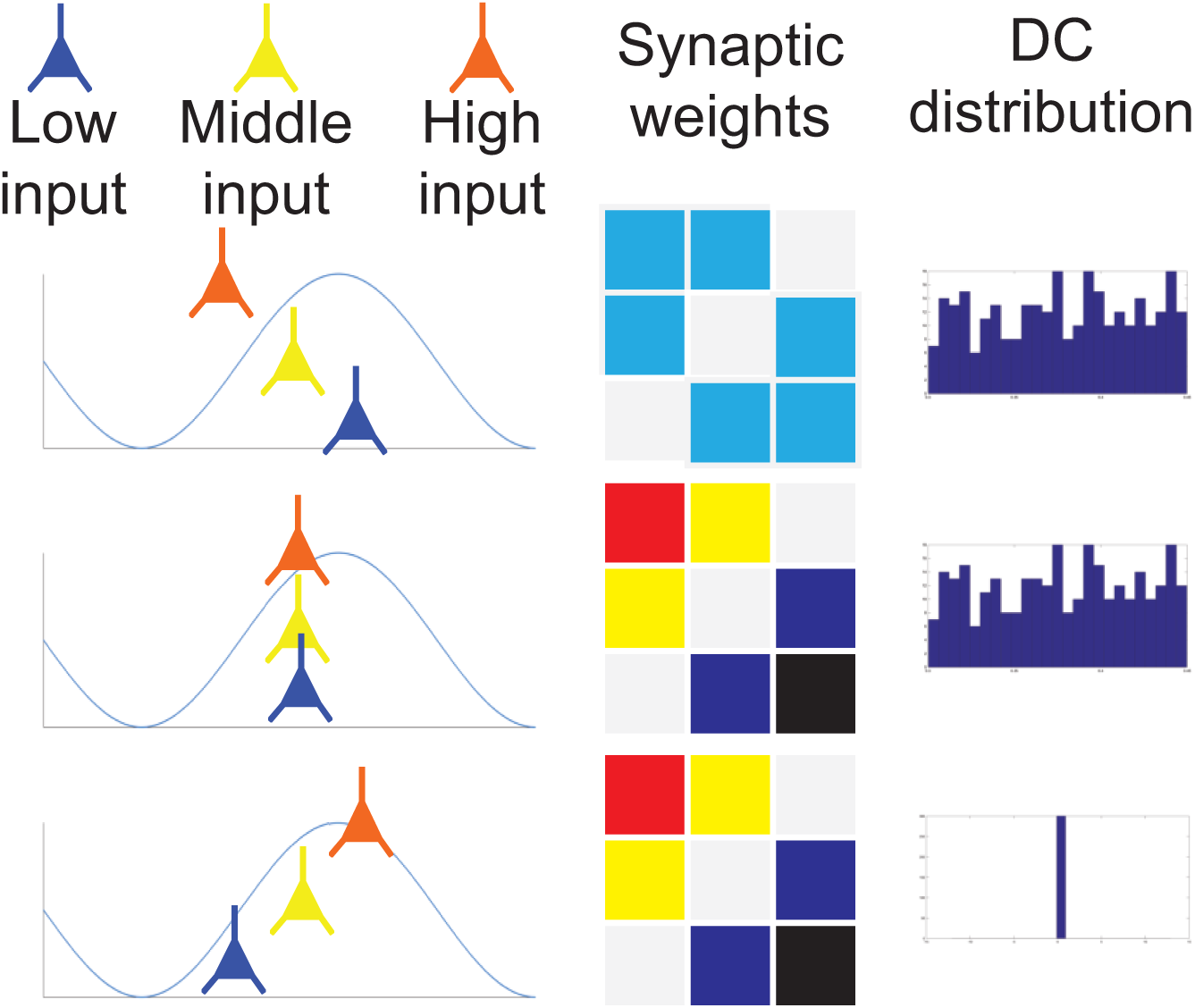
The model purposes a mechanism for the generation of reverse replay. Reverse replay due to how an input pattern imposes a phase procession of neuron firing due with respect to the oscillation. As the network learns the pattern inputs to weakly excited neurons are strengthened while those to highly excited cells are weakened. These synaptic changes compensate for the differences in excitation, leading to synchronous firing. When the pattern is removed the reverse mapping of synaptic weights leads to reverse reactivation.

Learning through STDP, or any Hebbian mechanism, requires either saturation or compensatory plasticity mechanisms to counteract the inherent positive feedback effects on firing rate, leading to network instability. Previous implementations of STDP have employed boundaries on synaptic weights, dynamic asymmetries between potentiation and depression, or renormalization of synaptic weights to preserve firing rates (reviewed in [27]). Our model proposes an alternative mode for preventing instability (Figure 5). As the input pattern is encoded into synaptic weights and the firing phase distribution becomes more uniform changes in synaptic weights decrease and stop due to features of the STDP curve around Δ*t* = 0, which is a reasonable fit to experimental data [26]. While many plasticity mechanisms exist both at the cellular and network level, the current mechanism provides an elegant solution to the question of when neural networks terminate learning of input patterns.

We have shown experimentally that our model agrees with pattern formation observed in hippocampal networks when channelrhodopsin-expressing PV+ interneurons are rhythmically stimulated. Within the hippocampus, functional network structure emerges or stabilizes during stimulation in the theta band (4-10 Hz). Using several methods of measuring functional connectivity within networks, we found a robust resonance effect in the formation of stable network structure (Figure 6). This effect is due to the organizing the firing of the network around the phase of the oscillatory input. That this effect is reproducible in various neuronal models [20] and *in vivo* suggests that it may be a general feature of activity is organized in neural networks, to optimize encoding of input patterns.

The input-dependent organization of network activity facilitated by subthreshold resonance provides a network-level substrate for sequential learning (Figure 5). When subsets of neurons have overlapping activation curves the relationship between input and firing phase creates spike time differences that are optimized for encoding the sequence order. One requirement for this result is that the activation of neurons needs to be skewed in time - in other words, repolarization occurs more rapidly than depolarization (Figure 5A). This ensures that connections strengthened by a balanced STDP regime are feedforward with respect to the sequence order, while feedback connections are weakened. Within the context of hippocampal place cells sequences, there is some evidence for this required skewness in activation [28, 29], though in an experience dependent manner [30]. Replay is the most direct readout of sequential learning. In the hippocampus, replay of place cell sequences occur both in the forward and reverse direction [16, 28, 17, 19]. These replay modes are represented in different proportions across behavioral states, with reverse replay being more prevalent during sleep [19]. In our model, forward replay occurs when a network is driven by noise (i.e. randomly activated) and reverse occurs when the network is reactivated by oscillating input (Figure 5D-F). These two network-activation states capture aspects of the awake resting (no theta rhythm, and decreased sensory activation, leading to forward replay) and sleep (theta presence, and low or decreased sensory activation, leading to reverse replay).

Hippocampal place cells show a theta phase precession in their firing, as an animal approaches a location neurons which code for a near-by place will fire in the troughs of the theta oscillation while those which code for a far place fire near the peak [14]. This phenomenon has also been shown in the entorhinal cortex [13] and in the ventral striatum ([15]. In our model cells in resonance with an oscillating rhythm they show a similar firing versus phase relationship.

Beyond the context of place cells, our model demonstrates how a network can translate information between the two main modes of neural coding rate [31] and phase [32, 33, 34, 35] coding. Both rate coding, where stimuli are represented by the firing rate of neurons, and phase coding, where information is represented in the time differences between spikes, are observed in nervous systems. Rate coding is a simpler and due to its super-threshold nature more reliable it is limited in its functionality in dynamic pattern separation [36]. Our results provide a mechanism for the translation between these two coding schemes and allows for networks to switch through neuromodulation [21]. While the extent to which this precise mechanism is responsible for information encoding in the brain remains an open question, our present data suggest that it has explanatory value for many of the observed *in vivo* phenomena surrounding learning.

